# EEG-based auditory attention detection: boundary conditions for background noise and speaker positions

**DOI:** 10.1101/312827

**Authors:** Neetha Das, Alexander Bertrand, Tom Francart

## Abstract

**Objective:** A listener’s neural responses can be decoded to identify the speaker the person is attending to in a cocktail party environment. Such auditory attention detection methods have the potential to provide noise suppression algorithms in hearing devices with information about the listener’s attention. A challenge is the effect of noise and other acoustic conditions that can reduce the attention detection accuracy. Specifically, noise can impact the ability of the person to segregate the sound sources and perform selective attention, as well as the external signal processing necessary to decode the attention effectively. The aim of this work is to systematically analyze the effect of noise level and speaker position on attention decoding accuracy.

**Approach:** 28 subjects participated in the experiment. Auditory stimuli consisted of stories narrated by different speakers from 2 different locations, along with surrounding multi-talker background babble. EEG signals of the subjects were recorded while they focused on one story and ignored the other. The strength of the babble noise as well as the spatial separation between the two speakers were varied between presentations. Spatio-temporal decoders were trained for each subject, and applied to decode attention of the subjects from every 30s segment of data. Behavioral speech recognition thresholds were obtained for the different speaker separations.

**Main results:** Both the background noise level and the angular separation between speakers affected attention decoding accuracy. Remarkably, attention decoding performance was seen to increase with the inclusion of moderate background noise (versus no noise), while across the different noise conditions performance dropped significantly with increasing noise level. We also observed that decoding accuracy improved with increasing speaker separation, exhibiting the advantage of spatial release from masking. Furthermore, the effect of speaker separation on the decoding accuracy became stronger when the background noise level increased. A significant correlation between speech intelligibility and attention decoding accuracy was found across conditions.

**Significance:** This work shows how the background noise level and relative positions of competing talkers impact attention decoding accuracy. It indicates in which circumstances a neuro-steered noise suppression system may need to operate, in function of acoustic conditions. It also indicates the boundary conditions for the operation of EEG-based attention detection systems in neuro-steered hearing prostheses.

**Index Terms:** Auditory attention detection, EEG processing, neuro-steered auditory prostheses, brain-computer interface, cocktail party, acoustic conditions.

The work is funded by KU Leuven Special Research Fund C14/16/057 and OT/14/119, FWO project nrs. 1.5.123.16N and G0A4918N, the ERC (637424) under the European Union’s Horizon 2020 research and innovation programme, and a research gift of Starkey Hearing Technologies. The scientific responsibility is assumed by its authors.

## 1. INTRODUCTION

Hearing aids have noise reduction algorithms that help improve speech perception by reducing undesirable noise. They however lack an important piece of information - the intent of the user. A multi-speaker environment with a lot of background noise is a challenging situation for a hearing impaired person who wants to listen to one speaker, and ignore the rest of the auditory stimuli present. Although a hearing aid can help reduce the surrounding noise, most hearing aids are not ‘smart’ enough to work in synchrony with the user’s intent. Hearing devices equipped with multiple microphones are capable of using beamforming techniques to improve the signal-to-noise ratio (SNR) in a given direction, but this information about the desired direction is not available.

It is well established that a listener’s cortical responses entrain to the envelopes of the speech streams present, and that the entrainment is stronger for the attended speech stream (Ding and Simon, 2012a; Golumbic et al., 2013). Auditory attention detection (AAD) algorithms allow to decode neural responses to auditory stimuli to detect the speaker a person is attending to, based on the auditory sources present, and the electroencephalogram (EEG) or magnetoencephalogram (MEG) of the person. Recent years saw progress in this area such as attention detection based on forward and backward modeling using various speech representations such as envelopes (O’Sullivan et al., 2014), spectrograms, phonetic features etc (Di Liberto et al., 2015), addressing issues such as the number of electrodes (Mirkovic et al., 2015), electrode placements (Fiedler et al., 2016; Mirkovic et al., 2016), the length of the data window to decode attention (Mirkovic et al., 2015; Zink et al., 2017), the use of concealable miniature devices (Fiedler et al., 2016; Mirkovic et al., 2016; Mundanad Narayanan and Bertrand, 2018), auditory-inspired envelopes (Biesmans et al., 2017) etc. We also saw the application of neural network based approaches to decode attention (Taillez et al., 2017; Wong et al., 2018a). Attention decoding has also been explored using canonical correlation analysis based decoders (de Cheveigné et al., 2018) as well as state-space estimators of attentional state rather than the more often used correlation measures (Akram et al., 2016; Miran et al., 2018). These developments bring us closer towards achieving the goal of faster attention detection which is a necessity if neuro-steered hearing aids are to become a reality. In addition, there has been some work towards building an AAD integrated noise suppression system (Das et al., 2017; O’Sullivan et al., 2017; Van Eyndhoven et al., 2017).

While work on algorithms for attention decoding and its integration into noise-suppression systems advance, there is a need for insights into their performance in realistic auditory scenarios. From the work by Das et al. (2017), we have seen that from an audio signal processing point of view, good attention decoding accuracies, as well as output signal-to-noise ratios are possible for a range of background noise levels and speaker positions. However, Das et al. (2017) used EEG data from recordings where the stimuli did not include background noise and with a maximal spacing between the two competing speakers, and hence the outcome could be considered rather optimistic. Since we rely on the SNR of the neural responses to perform attention decoding, it is important to have a systematic understanding of the impact of changing noise level and speaker positions on the quality of these neural responses. The boundary conditions for factors like noise level and speaker separation necessary for the operation of attention detection systems are still unknown. Fuglsang et al. (2017) analyzed the effect of reverberation and number of competing talkers, showing that cortical activity tracked the attended speech stream and the tracking remained robust irrespective of the realistic acoustic environments used in the study. It was also found that more realistic stimuli (filtered by head-related transfer functions) result in better attention decoding performance compared to dichotically presented stimuli (Das et al., 2016). There has also been research into how attention decoding is impacted when unprocessed binaural signals are used as reference signals in different acoustic environments (reverberant and/or noisy) (Aroudi and Doclo, 2017).

In this study, we focus on the impact of background babble noise as well as relative separation between the two competing talkers, on AAD. We used negative signal to noise ratios that can be expected to result in significantly different speech intelligibilities, as well as relative speaker positions which can be perceived to be easy to hard in terms of speech intelligibility and the effort necessary to focus on one speaker and ignore the other. Finally, drawing inspiration from the work of Vanthornhout et al. (2018), we looked into the relationship between speech intelligibility and attention decoding accuracies.

## II. METHODS

### A. Data Acquisition

Twenty-two female and six male subjects between the ages of 20 and 25 years participated in the experiment. All subjects were native Dutch speakers. Pure-tone audiometry was performed by which all subjects were found to be normal-hearing (thresholds < 20 dB HL from 125 Hz to 8000 Hz). Every subject signed an informed consent form approved by the local ethical committee.

During the experiment, 64-channel EEG data at a sampling rate of 8192 Hz was recorded using the BioSemi ActiveTwo system. The speech stimuli (competing talkers) were administered to the subject at 65 dBA through Etymotic ER1 insert earphones. The experiments were conducted in a soundproof, electromagnetically shielded room. We used the APEX 3 program developed at ExpORL (Francart et al., 2008) to present sounds and an interactive screen to the subjects where they could use a mouse to navigate through the experiment.

### B. Stimuli and experiment design

We chose Dutch short stories narrated by male speakers as stimuli. In the audio files, silences longer than 300 ms were shortened to 300 ms. For each story, the root mean square (RMS) value of the speech material was computed for every 0.5 second frame, which formed the vector of RMS values. The RMS intensity over time of the stories was smoothed so that there were no large variations in the sound pressure level of the stimuli over time. This was done to avoid differences in speech intelligibility and also to prevent a shift in attention of the subject if and when significant sound pressure level differences occur between the attended and the unattended speakers. All stories were then normalized to the same overall RMS intensity.

We made 9 babble noise files by combining speech streams from 2 unique male and 2 unique female speakers at a time. Head-related transfer functions (HRTFs) (Kayser et al., 2009) were used to filter each of these babble noise streams such that babble noise was perceived to come from 9 equidistant (from -180 degrees to 140 degrees in steps of 40 degrees) locations around the subject. The long-term spectrum of the babble noises was matched to the average spectrum of all the stories used in the experiments. The overall effect to the subject was that of a 36-talker (9 directions X 4 speakers) surrounding babble. We define an experiment here as a sequence of 8 presentations. An example for the design of an experiment is shown in Table I. Within an experiment, each presentation consisted of the subject being presented with two story parts at 65 dBA - each from a particular direction with respect to the subject, as well as surrounding babble noise at a particular signal to noise ratio (SNR) with respect to the power of the speakers. SNR in this work corresponds to the ratio of signal power of one of the competing talkers to the total signal power of the 9 babble noise streams. The screen in front of the subjects showed a figure with indication towards the direction of the speech stream to which the subjects were expected to focus their attention. Each subject participated in 4 experiments where the speaker positions were at -90 (extreme left) and 90 (extreme right) degrees, 30 and 90 degrees, -30 and -90 degrees, and -5 and 5 degrees. Thus, the design allowed to analyze situations where the angular separation between the speakers were 180 degrees, 60 degrees and 10 degrees respectively.

**TABLE I:**
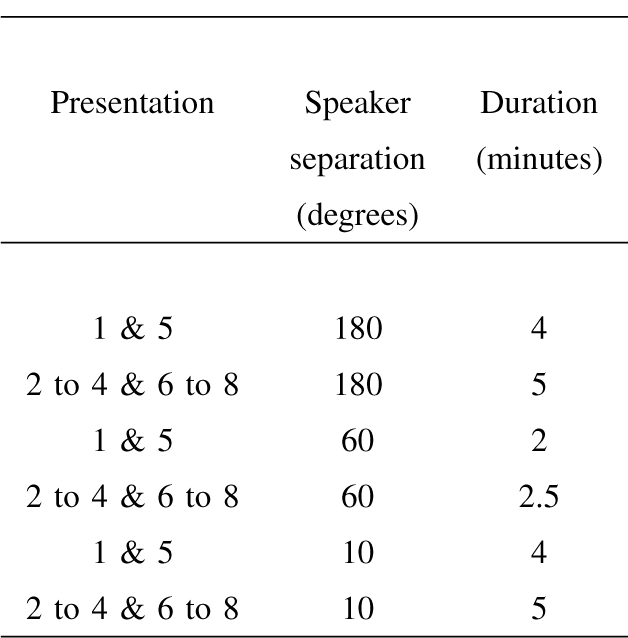
Duration of each presentation for different angular separations between speakers.

Before the first presentation, the subject was presented with a test screen and a 5 s audio of only the attended speaker so that the subject could get an idea of the attended speaker’s voice and direction before the presentations began. In the first presentation, the two speakers were presented to the subject with no background babble. In the following sections, we will be referring to this as the ‘no noise’ condition. For convenience, the ‘no noise’ condition is shown as 20 dB SNR in all the tables and figures. In the following 3 presentations, the speakers were presented along with surrounding babble at 3 different SNRs: -1.1 dB, -4.1 dB and -7.1 dB. After the first 4 presentations, the subject was again presented with a test screen, this time accompanied by indication and a 5 s audio of the other speaker. For presentations 5 to 8, the subject was instructed to focus his attention on the other speaker. The order of SNRs, the presented stories and the order of attended direction were randomized between experiments as well as subjects to avoid any possible bias. After each presentation, the subjects had to answer a multiple choice question about the content of the attended story. The subjects also had to mark on a scale on the screen, what percentage of words of the attended speaker they could understand. After each experiment, we also repeated the presentations 1 and 5 so that we could collect enough data in the ‘no noise’ condition to independently train decoders on data with no exposure to babble noise, if the need arose. Table II shows the duration of presentations for different speaker separations.

**TABLE II:**
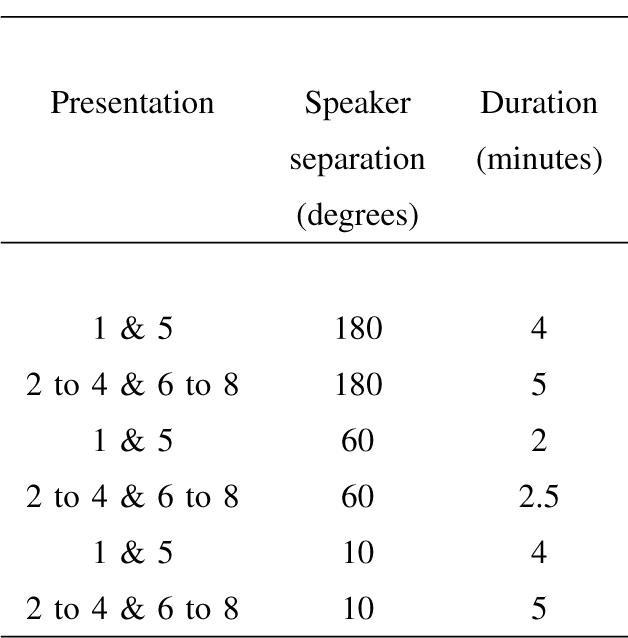
An example of the design of an experiment where speech streams were presented from -5 degrees (left) and 5 degrees (right) with respect to the frontal direction of the subject.

#### Choice of SNRs

We chose signal to noise ratios that would ensure significantly different speech intelligibility levels. For this, we conducted an adaptive procedure to find the speech recognition threshold (SRT), i.e. the SNR at which speech intelligibility was 50%. The subject had to attend to a list of ‘Matrix’ sentences (Luts et al., 2014), in the presence of a competing talker, and surrounding babble noise for the 3 speaker separations. The SNR was changed adaptively (Brand and Kollmeier, 2002) until the point where the subject could recognize 50% of the words in the sentence correctly. This behavioral test was done before the start of the main experiment. Thus, standard SRTs for the 3 speaker separations were found for 13 out of the 28 subjects who participated. (The first 15 subjects performed the test for only 180 degree speaker separation, and hence the corresponding data was incomplete for analyzing the correlation between attention decoding performance and speech intelligibility.) An experiment was also designed to find subjective SRTs in the case of non-standardized speech material such as the short stories used in our main experiment (Decruy et al., 2018). During this adaptive procedure, subjects were asked to indicate on the screen whether they understood more or less than 50% of the attended story or sentence. The SNRs were adapted accordingly until the subjects reached the 50% speech intelligibility point. Subjective SRTs found using this method (‘rate’ method) as well as standard SRTs found using the standard adaptive procedure (‘recall’ method) were compared for the Matrix speech material. On a set of 8 subjects who participated in a pilot study, the results for the ‘rate’ and the ‘recall’ methods using the Matrix speech material differed, for the 3 speaker separations (figure 1). We found a difference of 1.3 dB between the mean ‘rate’ (subjective) and ‘recall’ (standard) SRT, for the 180 degree speaker separation for the Matrix speech material. Since standard tests can not be used for finding the SRT for story speech material, we applied the 1.3 dB as a correction on the mean ‘rate’ SRT for the story speech material to find an estimated standard SRT for story speech material. At this estimated SRT (-7.1 dB), we could expect 50% intelligibility for the story speech material for the speaker separation of 180 degrees. This was chosen to be the lowest SNR for the main experiment. We then chose 2 more SNRs -4.1 dB and -1.1 dB, increasing the first chosen SNR in steps of 3 dB, similar to the method of choosing SNRs by Petersen et al. (2016). We, however, refrained from using individualized SNR levels as done by Petersen et al. (2016), since we tested only normal hearing subjects. In addition to these 3 SNRs, we also recorded data with no babble noise in the background.

**Fig. 1:**
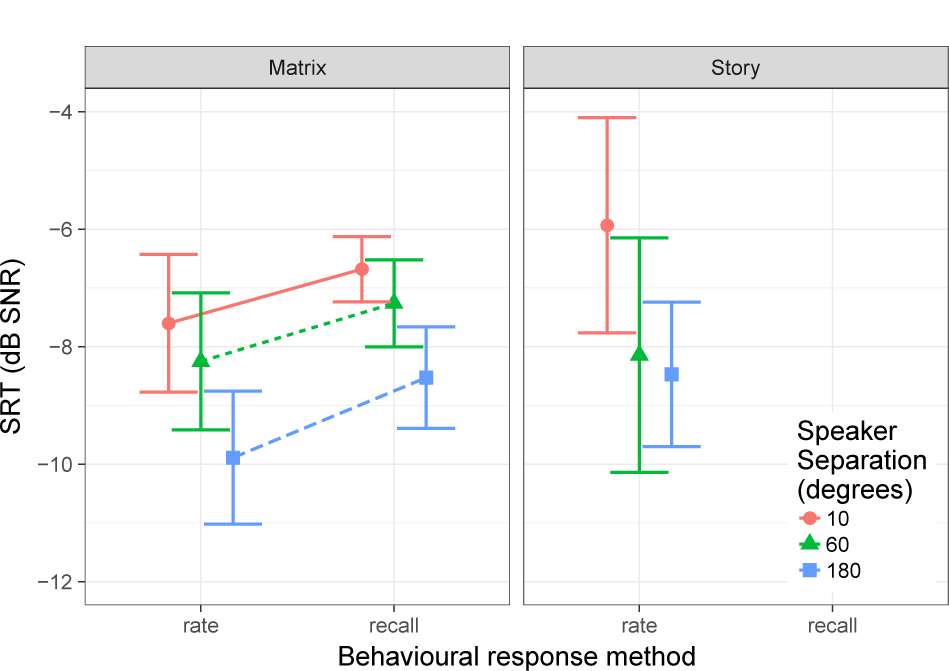
Speech recognition thresholds for different speech material. Offset between the mean SRTs of the ‘recall’ and ‘rate’ methods using Matrix speech material is used as a correction for the mean ‘rate’ SRT when story speech material is used. Lower SRT corresponds to better speech intelligibility.

#### Choice of speaker positions

Speaker positions were chosen to be representative of different real-life situations. We chose 180 degree speaker separation to represent an intuitively easy scenario, where spatial release from masking would make auditory object formation and selection an easy task. We also included the speaker positions -5 and 5 degrees which is an extreme case in which one can imagine two speakers standing shoulder to shoulder at a distance of 3 meters in front of the subject. Compared to the 180 degree condition, this scenario can be expected to be difficult since the speakers are almost co-located, and a certain amount of effort would be necessary in order to attend to one speaker and ignore the other. This is confirmed by the SRTs in 180 and 10 degree speaker separations shown in figure 1 for the ‘recall’ method using Matrix speech material. These conditions are similar to the test conditions in the LiSN-S test (Cameron and Dillon, 2007). The third choice for speaker separation was 60 degrees, in which case, both the speakers were on the same side of the head. Das et al. (2017) found that in a system performing neuro-steered noise suppression at low input SNRs, speaker positions on the same side of the head resulted in a relatively lower improvement in output SNRs compared to symmetric speaker set ups^1^. To balance between the two sides of the head, to avoid introducing a lateralization bias during the training of decoders (Das et al., 2016), we split this condition between two experiments where the speakers were at -30 and -90 degrees, and 30 and 90 degrees. During our analysis, we combined the results of these two experiments, and only analyzed the differences between the three speaker separations: 10 degrees, 60 degrees and 180 degrees.

#### C. Attention decoding

##### 1) Data Preprocessing

The recorded EEG data was bandpass filtered between 0.5 and 10 Hz, using an equiripple bandpass filter with a maximal pass band attenuation of 0.5 dB, transition width of 0.9 Hz, and stop band attenuations of 20 dB (lower stop band) and 15 dB (upper stop band). This choice of the frequency range was based on earlier studies (Das et al., 2016; Ding and Simon, 2012b; Golumbic et al., 2013; Pasley et al., 2012) that showed the best attention decoding performance based on cortical envelope tracking in this range. The filtered signal was then downsampled to 40 Hz. For the story stimuli presented to the subjects, a gammatone filterbank was applied to split the speech signal into 15 filter bands, and a powerlaw compression with exponent 0.6 was then applied to each of these subbands (Patterson et al., 1995). The subbands were bandpass filtered using the same equiripple filter used for the EEG signals, and finally they were combined with equal weights, and downsampled to 40 Hz, to form a single ‘powerlaw subband’ envelope per stimulus. Powerlaw compression helps to replicate the non-linear transformation of the stimulus with relatively higher attenuation of higher amplitude signals, similar to the human auditory system (Stevens, 1955). Biesmans et al. (2017) found that this method of auditory inspired envelope extraction resulted in significantly better attention decoding accuracies than other envelope extraction methods. The data set was divided into 30 second trials which resulted in 276 trials per subject, equally divided between the three speaker separations.

##### 2) Stimulus Reconstruction and Decoder Design

We used the algorithm described by Biesmans et al. (2017) for our decoder design. ‘Powerlaw subband’ envelopes *s*_*a*_(*t*) and *s*_*u*_(*t*) were extracted from the attended and unattended speech signals respectively. A linear spatio-temporal decoder was used to reconstruct the envelope of the attended speech signal from the EEG data. A decoder consists of weights *D*_*c*_(*n*) for each EEG channel *c* and each time lag *n*. The time lags represent the difference between the time of stimulus presentation and the time when this stimulus is reflected in the cortical response. We used time lags of up to 250ms (O’Sullivan et al., 2014). If EEG data of channel *c* is represented by *M*_*c*_(*t*) where *t* is the time index, the reconstructed envelope on the attended speech signal is given by:

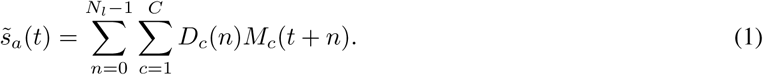

where *C* denotes the number of channels and *N*_*l*_ denoted the number of time-lags used in the decoding. For the sake of notation, we stack all the *D*_*c*_(*n*) in a vector 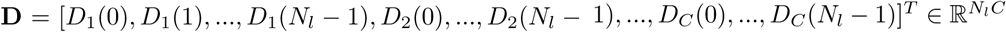, and we concatenate *M*_*c*_(*t*) and lags in a vector 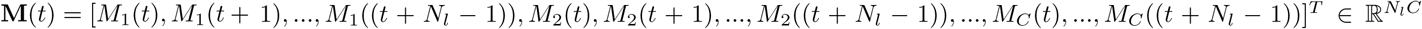. The design of the decoder is in such a way as to minimize the mean squared error between the attended speech envelope *s*_*a*_(*t*) and its reconstruction 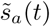. The decoder for each trial is found based on all other trials by solving the standard minimum mean squared error problem:

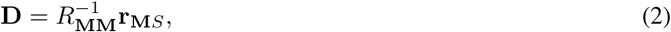

where 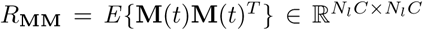 is the correlation matrix of the EEG data which includes cross correlation values between all combinations of channels and lags, and 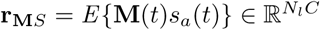 is the cross-correlation vector between the attended speech envelope and the EEG data, over all channels and time lags (where *E*{.} denotes the expectation operator).

For each subject, we used 30 s trials (i.e. *L* = 30 s x 40 Hz = 1200 samples) for decoding attention. Thus the correlation matrix for the *k*^*th*^ trial can be found by first concatenating **M**(*t*) at each time index as columns of the matrix 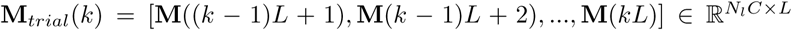, and then computing 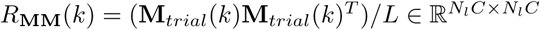. Similarly, the cross-correlation vector for the *k*^*th*^ trial can be found by first taking the attended speech envelope **s_a_**(*k*) = [*s*_*a*_(*k -* 1)*L* + 1)*, s*_*a*_(*k -* 1)*L* + 2)*, …, s*_*a*_(*kL*)]^*T*^ *∈*ℝ^*L*^ and then computing 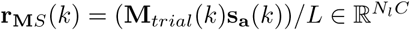.

Adopting a leave-one-trial-out approach for training decoders, the correlation matrices and the cross-correlation vectors are estimated over the samples from all trials except the test trial. As done by Biesmans et al. (2017), these correlation matrices and cross correlation vectors are averaged to find a mean correlation matrix and a mean cross-correlation vector. This process of averaging equates to concatenating all the data and then finding a correlation matrix and cross correlation vector, and results in better conditioned correlation matrices. The mean correlation matrix and mean cross-correlation vector are then used to compute the decoder from equation 2, after which it is applied to the test trial’s EEG data to obtain the reconstructed envelope. By opting for this method, instead of finding a decoder on a per-trial basis and then averaging the decoders (as originally proposed by O’Sullivan et al. (2014)), we eliminate the need to use regularization schemes (provided sufficient amount of data is available) such as ridge regression, Tikhonov regularization (first derivative type) etc (Wong et al., 2018b).

In this manner, a reconstructed attended envelope 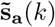 was found per trial, for which Pearson’s correlation coefficient was then computed with that trial’s **s_a_**(*k*) and **s_u_**(*k*). Attention was considered to be decoded correctly if the correlation of 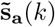 with **s_a_**(*k*) was found to be higher than with **s_u_**(*k*). These results were then grouped based on stimuli conditions and averaged across trials to obtain percentage decoding accuracies per subject for these conditions.

## III. RESULTS

### A. Mean attended and unattended correlations

Attended and unattended correlations were calculated for each trial, and we analyzed the influence of SNR and speaker separation on the mean correlations. Remarkably, as can be seen in the figure 2, we observe that both mean attended and mean unattended correlations were lower in the ‘no noise’ condition (to the right of the vertical lines) compared to when some noise is added. However, within the noisy conditions, we observed that mean attended correlations had a tendency to drop linearly with increasing noise power. Therefore we excluded the ‘no noise’ condition data while fitting linear mixed effect models for the correlations. We used the software R (version 3.4.3), and the R package ‘nlme version 3.1-137’ (Pinheiro et al., 2018) for fitting the linear mixed effect models. All the linear mixed effect models in this work were fitted by maximizing the restricted log-likelihood, and the residuals were checked for normality to ensure a good fit. We fitted 2 linear mixed effect models: one which models attended correlations as a function of SNR and speaker separation, and their interaction (fixed factors), and subject as a random factor, and the second which models unattended correlations as a function of the same fixed factors and subject as the random factor. We found that while mean attended correlations dropped significantly (*F*_221_ = 96.286*, p <* 0.001) with lower SNRs, SNR did not have a significant influence on mean unattended correlations. We also found that speaker separation significantly influenced both mean attended (*F*_221_ = 65.434*, p <* 0.001) and mean unattended correlations (*F*_221_ = 64.715*, p <* 0.001). In addition, we found a significant interaction effect of SNR and speaker separation (*F*_221_ = 5.773*, p* = 0.017) on mean attended correlations. Post-hoc Tukey tests with Bonferroni correction were performed to compare between groups of SNR and speaker separation. We found that mean attended correlations at -7.1 dB SNR were significantly lower than those at -1.1 dB (*z* = 6.775*, p <* 0.001) and -4.1 dB (*z* = 4.411*, p <* 0.001). The mean attended correlations at 180 degrees speaker separation were found to be significantly higher than at 60 degrees (*z* = 4.424*, p <* 0.001) and 10 degrees (*z* = 6.265*, p <* 0.001). The post-hoc tests on mean unattended correlations revealed that the unattended correlations dropped as the speaker separation increased (unattended correlations at: 10 degrees > 60 degrees with *z* = –2.977*, p* = 0.009, 60 degrees > 180 degrees with *z* = –3.098*, p* = 0.005, and 10 degrees > 180 degrees with *z* = –6.074*, p <* 0.001).

**Fig. 2:**
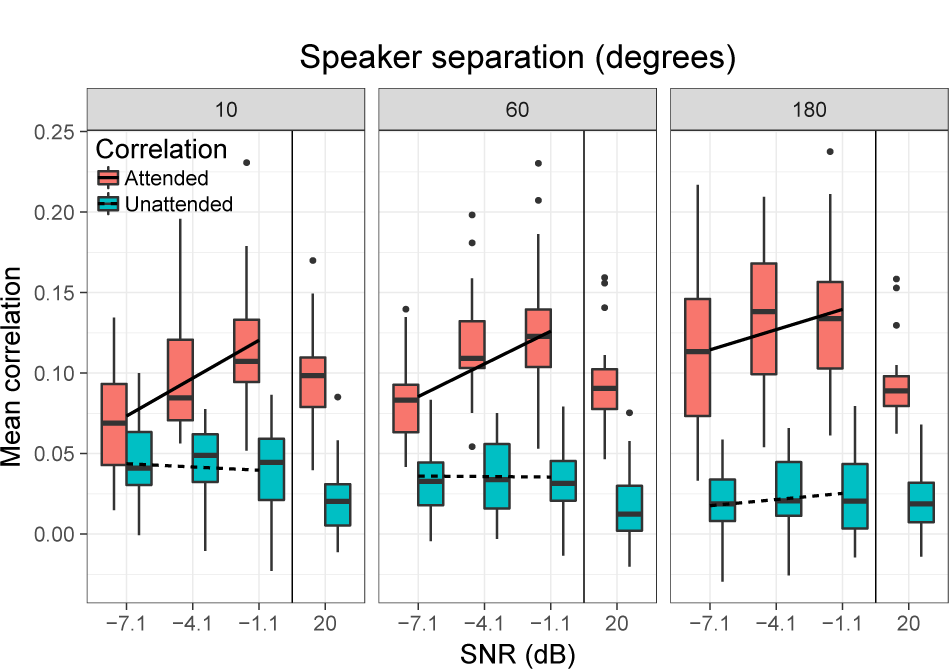
Boxplots showing mean attended and unattended correlations across subjects for different SNRs and speaker separations. The horizontal lines show predictions of the linear mixed effect models for each speaker separation.

### B. Attention decoding accuracy

Attention decoding accuracy was calculated based on trials grouped with respect to the different auditory conditions in the experiments. Figure 3 shows mean decoding accuracy (over all 28 subjects) for the different SNRs and speaker separations. Thus, we analyzed the performance differences across the 4 SNR conditions: -7.1 dB, -4.1 dB, -1.1 dB and 20 dB, as well as across the 3 speaker separations: 10, 60 and 180 degrees.

**Fig. 3:**
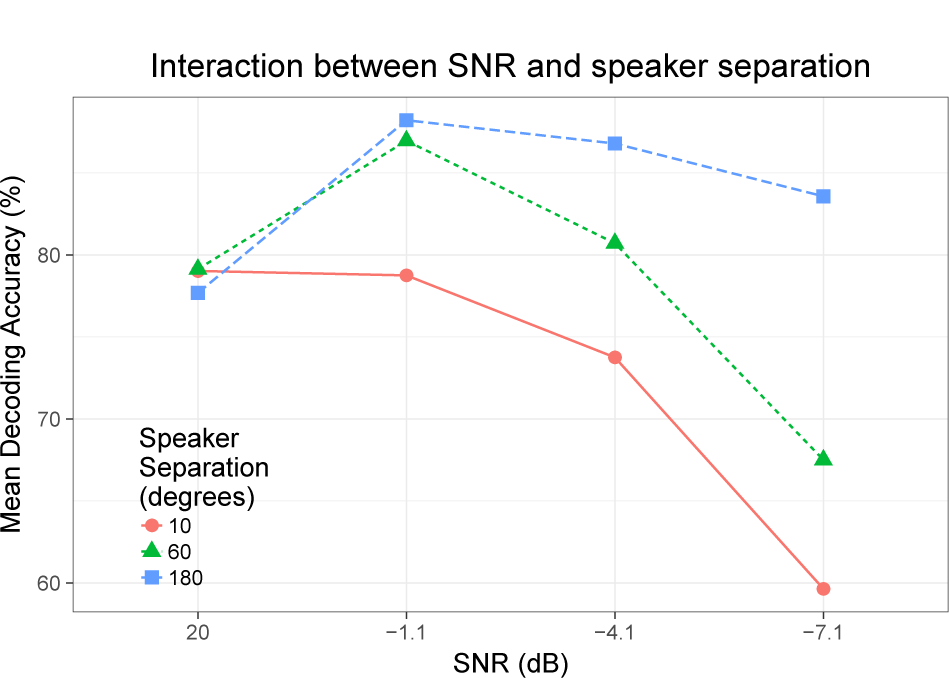
Mean decoding accuracies for the 4 SNRs and 3 speaker separations showing interaction between noise power and the angular separation between the competing talkers.

To investigate the overall effect of SNR and speaker separation on attention decoding accuracy, a linear mixed effect model was used to model the relationship between mean decoding accuracy and SNR, speaker separation and their interaction, with subject as random factor. The linear mixed effect model was fitted using all the data excluding the ‘no noise’ condition. The results showed that the attention decoding accuracies dropped significantly with SNR (*F*_221_ = 68.415*, p <* 0.001) as well as speaker separation (*F*_221_ = 74.904*, p <* 0.001) (figure 3). We also found a significant interaction effect of SNR and speaker separation (*F*_221_ = 14.242*, p <* 0.001). Post-hoc comparisons (Tukey test with Bonferroni correction) between decoding accuracies grouped by SNR revealed that decoding accuracies at -7.1 dB were significant worse compared to those at -4.1 dB (*z* = 4.737*, p <* 0.001) and at -1.1 dB (*z* = 6.416*, p <* 0.001). The relationship between decoding accuracy and SNR can be observed in figure 4.

**Fig. 4:**
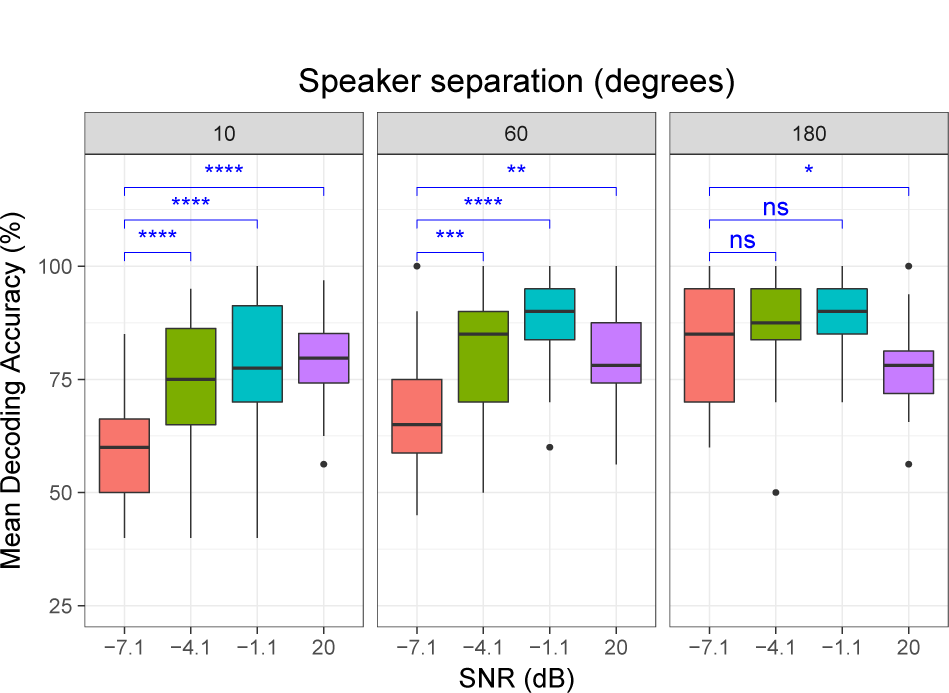
Decoding accuracies based on trials grouped by SNRs.

We also analyzed the impact of SNR on attention decoding accuracy for each of the speaker separations. We performed paired t-tests (with Bonferroni correction) between decoding accuracies grouped by SNR for each speaker separation (figure 4). The significance level for differences between different groups are indicated by: ‘*’ for *p <* 0.050, ‘**’ for *p <* 0.010, ‘***’ and ‘****’ for *p <* 0.001. We found an interesting trend here: with respect to the ‘no noise’ condition, attention decoding accuracy increased when some noise was present for both 180 and 60 degree speaker separations. In the noisy conditions, as can be seen in figure 4, decoding accuracies dropped with SNR for the 10 and 60 degree speaker separations.

Post-hoc comparisons (Tukey tests with Bonferroni correction) were also done to look for differences in decoding accuracies grouped by speaker separation (figure 5). A comparison between groups revealed that decoding performance decreased with speaker separation (decoding accuracies at 10 degrees < 60 degrees with *z* = 2.638*, p* = 0.025, 60 degrees < 180 degrees with *z* = 5.397*, p <* 0.001, and 10 degrees < 180 degrees with *z* = 8.035*, p <* 0.001). The relationship between decoding performance and speaker separation can be observed in figure 5.

**Fig. 5:**
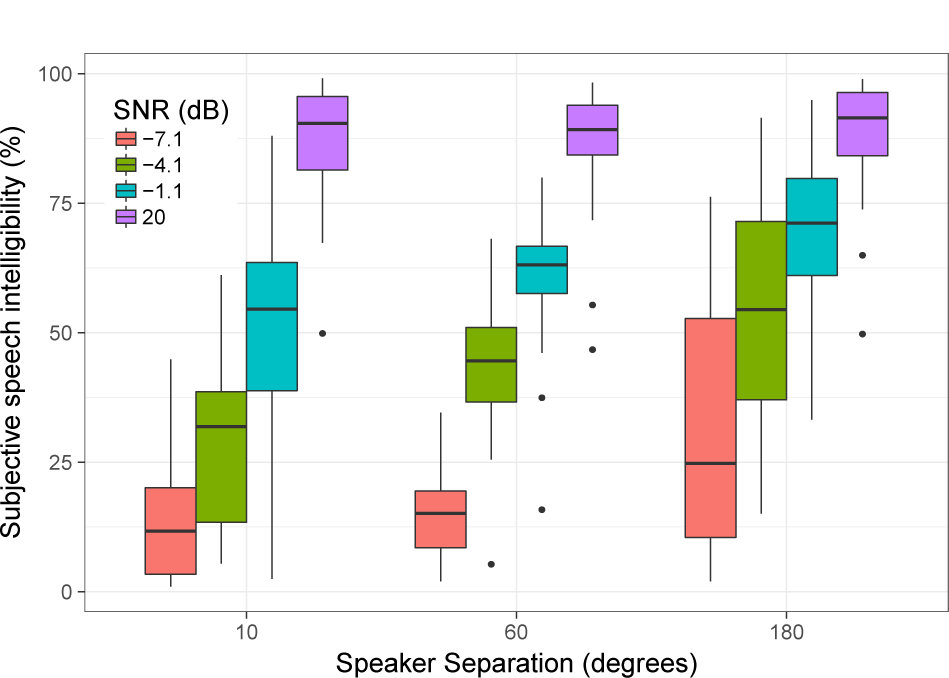
Decoding accuracies based on trials grouped by speaker separation.

We also performed paired t-tests (with Bonferroni correction) to investigate the effect of speaker separation on the decoding accuracies, for each of the SNR conditions. As can be seen in figure 5, for the ‘no noise’ condition, there was no significant effect of speaker separation on decoding accuracy. However, for the noisy conditions, it was seen that the decoding accuracy decreased with speaker separation.

### C. Comparison with speech intelligibility (SI)

Out of the 28 subjects who participated in the study, 13 subjects also participated in adaptive behavioral speech tests (‘recall’ method) to find their SRTs for the 3 speaker separations, using the Matrix speech material as the attended sentence and a story as the competing talker, with the power of the background babble adapted based on the subjects’ responses. From this data, we fitted psychometric curves for each subject and speaker separation to get speech intelligibility scores (for Matrix speech material) at the SNRs presented in the main experiments. For convenience we will refer to these scores as ‘standard’ speech intelligibility scores. We used a linear mixed effect model to investigate standard speech intelligibility with fixed factors as SNR, speaker separation and their interaction, and subject as the random factor. We found that standard speech intelligibility (for Matrix speech material) significantly dropped with SNR (*F*_96_ = 252.373*, p <* 0.001)as well as speaker separation (*F*_96_ = 28.608*, p <* 0.001). We also found a significant interaction effect of SNR and speaker separation (*F*_96_ = 10.001*, p <* 0.001). We calculated the Spearman correlation between attention decoding accuracy and standard speech intelligibility under the same conditions, and found a significant effect (*ρ* = 0.423*, p <* 0.001). We excluded the ‘no noise’ condition from the data for fitting the linear model because we found attention decoding accuracies to be significantly higher in the presence of moderate noise as compared to the ‘no noise’ condition.

During the experiments, all subjects reported their speech intelligibility (subjective) after each presentation. Figure 6 shows the mean of the speech intelligibility scores indicated by the 28 subjects grouped by the different SNR conditions and speaker separations. For convenience, we refer to this as ‘subjective’ speech intelligibility. We also investigated the relationship between subjective speech intelligibility and attention decoding accuracy. We fitted a linear mixed effect model for the subjective speech intelligibility with SNR, speaker separation and their interaction as fixed factors, and subject as random factor. We found that the subjective speech intelligibility dropped significantly with dropping SNR (*F*_216_ = 315.663*, p <* 0.001) as well as speaker separation (*F*_216_ = 36.326*, p <* 0.001).

**Fig. 6:**
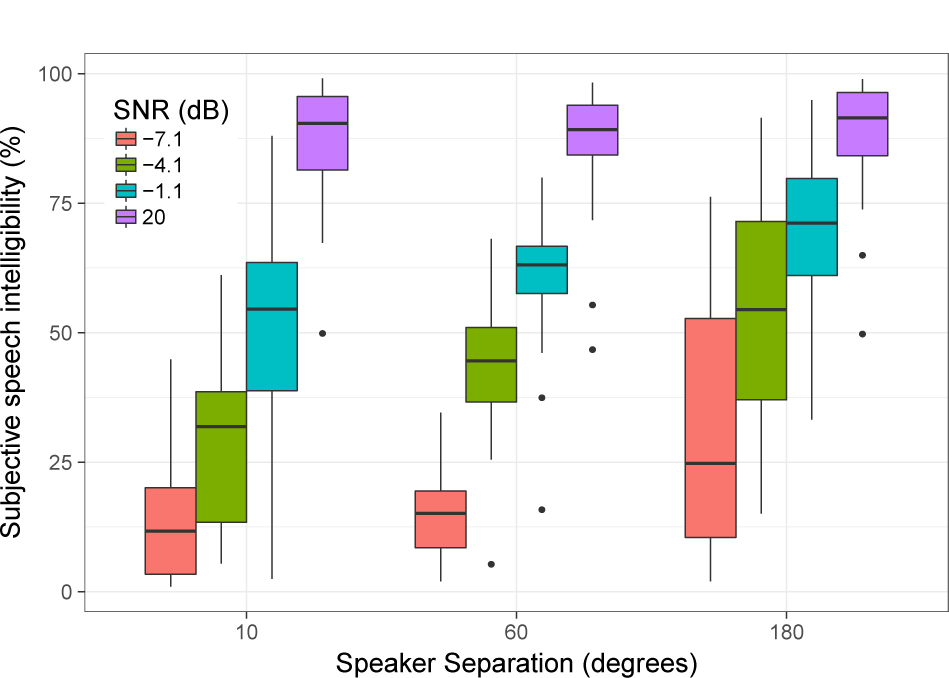
Reported speech intelligibility at different SNRs for each speaker separation.

Figure 7 shows how attention decoding accuracy varied with subjective speech intelligibility. We fitted a linear mixed effects model for attention decoding accuracy with the subjective speech intelligibility as fixed factor, and subject as random factor. The decoding accuracies from the ‘no noise’ condition were excluded from the model fitting. We found that attention decoding accuracy dropped significantly with subjective speech intelligibility (*p <* 0.001). The predicted values of the linear mixed effects model for attention decoding accuracy are plotted as a line in the figure 7. We also found a significant Spearman correlation (*ρ* = 0.591*, p <* 0.001) between attention decoding accuracy and subjective speech intelligibility.

**Fig. 7:**
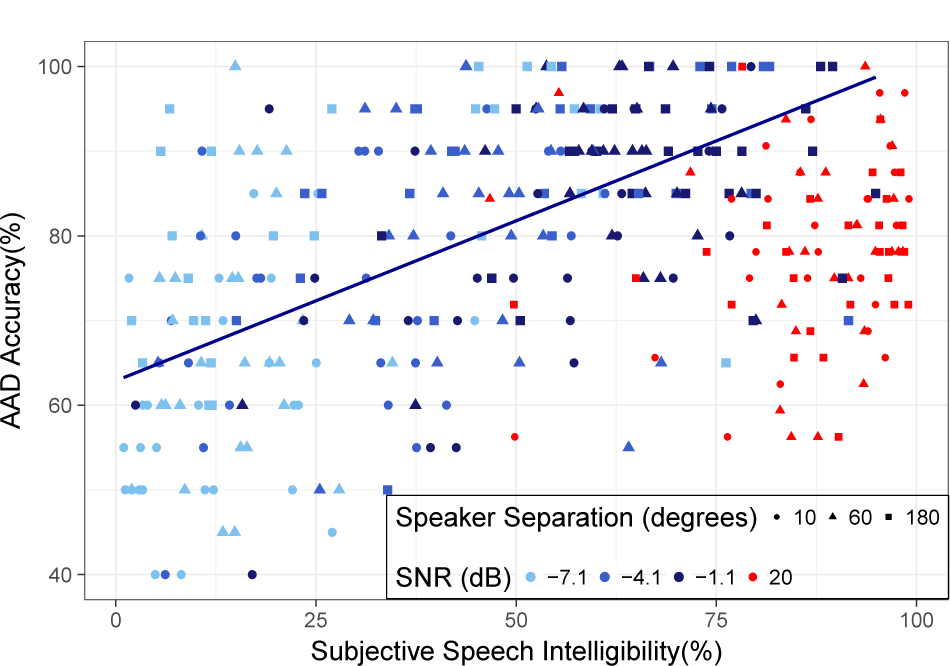
Attention decoding accuracies plotted against subjective speech intelligibilities of 28 subjects for the 4 SNRs and 3 angular separations between speakers. The points in red correspond to the ‘no noise’ condition. These points were excluded in the linear mixed model fitting (the dark blue line).

## IV. DISCUSSION

This study was designed to investigate the impact of challenging acoustic conditions on attention decoding performance. The chosen SNR values ensured significantly different speech intelligibility between conditions (figure 6). The mean attention decoding performance on 30s trials in the ‘no noise’ condition over all speaker separation conditions spanned a range between 59% and 97% which is similar to the attention decoding performance reported by Biesmans et al. (2017) and Das et al. (2016).

### A. Influence of noise power on attention decoding

We found that SNR adversely affects the attended speech representation in the neural responses, whereas the representation of the unattended speech envelope remains more or less unaffected. This is also in line with the study of Petersen et al. (2016) where the tracking of the attended speech was found to increase in magnitude with lower noise levels, while no such effect could be observed for the tracking of the ignored speech. Since we found that attended correlations dropped significantly with SNR, the separation between the attended and unattended correlations decreased with lower SNRs. This can be expected to have a direct influence on the attention decoding accuracy. Indeed, we found that attention decoding accuracy significantly drops with SNR.

Our analysis also shows how the interaction between background noise power and separation of speakers affect attention decoding accuracy (figure 3). We found that mean decoding performance was significantly lower in the ‘no noise’ condition compared to -1.1 dB SNR. With the inclusion of background noise, attention to one speaker becomes harder, and consequently demands more effort from the subject. The top-down processes involved in suppressing the additional noise could also result in stronger neural entrainment of the attended speech stream, and hence better signal-to-noise ratio in the EEG data. This could explain the relatively better decoding accuracies compared to the less challenging ‘no noise’ condition. However, as the background noise power increases, this benefit fails to outweigh the disadvantage of the noise. With rising noise power, sound segregation cues are degraded, and as a consequence, features extracted from the auditory scene can no longer be effectively grouped to represent a specific attended stream (Ding et al., 2014). This results in reduced cortical entrainment, and hence worse decoding performance as can be seen for -4.1 dB and -7.1 dB SNRs.

### B. Influence of speaker positions on attention decoding

From figure 5, it can be inferred that in challenging SNR conditions, decoding performance for smaller speaker separations is more severely affected than for speaker positions that are far apart (for eg. 180 degrees). In a cocktail party scenario, the listener performs sound segregation based on many features such as spectro-temporal cues, perceptual differences, and speaker locations. Binaural cues such as inter-aural time differences (ITD) and interaural level differences (ILD) occurring due to difference in spatial locations of the competing talkers are exploited to perform segregation of speech streams (Carlyon, 2004). Angular separation between speakers is expected to affect spatial release from masking (SRM). Kidd Jr et al. (1998) and Jones and Litovsky (2011) found the advantage of speaker separation in the identification of audio signal patterns, thereby improving speech intelligibility in a cocktail party scenario. In our study, therefore, the 10 degree speaker separation can be expected to be a more difficult auditory scene in terms of segregation of speech streams, in comparison to the wider speaker separations 60 and 180 degrees. This is in line with our findings that the 10 degrees speaker separation resulted in the worst SRT values from the behavioral tests. From figure 3, it can also be seen that the 10 degree speaker separation is indeed the least robust to background noise power, with decoding accuracies dropping more sharply than the other speaker separations as the noise power increases.

As opposed to SNR, speaker separation did have a significant effect on the correlation with the unattended speaker. Horton et al. (2013) describe an entrainment-based suppression mechanism that reduces correlations with the unattended speech stream during selective attention. In our study, the finding that unattended correlations increase with reducing speaker separation suggests that the suppression mechanism is adversely affected by noise for closer speaker separations. The difference between mean attended and unattended correlations was also found to decrease with decreasing speaker separation, which can be expected to result in reduction in attention decoding accuracy (figure 2).

### C. Relationship between speech intelligibility and attention decoding accuracy

We hypothesized that the top-down processes that support auditory object formation and selection, and suppression of noise to improving speech perception, are also responsible for the improvement in signal-to-noise ratio of the neural responses that support attention decoding under challenging auditory conditions. Zekveld et al. (2006) found brain regions involved in top-down processing supplementing speech comprehension to be more active when the speech was less intelligible or presented at low SNRs. Riecke et al. (2017) demonstrated that speech-brain entrainment contributes functionally to speech intelligibility, and that speech-entrainment and intelligibility interact reciprocally. Ding and Simon (2013) also found subjective speech intelligibility to be correlated with measures of neural synchronization with the stimulus. We therefore expected to see a significant correlation between the speech intelligibility reported by the subjects and their attention decoding accuracy. In figure 7, although we observed large variability in attention decoding accuracy with respect to subjective speech intelligibility, we indeed found a significant correlation between speech intelligibility and attention decoding accuracy. This is also supported by the findings of Vanthornhout et al. (2018) where the correlation between the reconstructed envelope from neural responses, and the acoustic speech envelope was proposed to be used as an objective measure of speech intelligibility. Rimmele et al. (2015) found more precise speech-tracking of attended speech to be related to processing its fine structure, possibly reflecting the application of higher-order linguistic processes. They did not find a similar dependence on the fine structure for the unattended speech. This, along with the dependence of attended correlations on the signal to noise ratio that we found in our study, suggests that the top-down processes that are involved in sound segregation and speech perception, thereby influencing speech intelligibility, could also be responsible for the cortical entrainment to the attended speech stream.

In figure 7, mean attention decoding accuracy can be seen to exceed 80% when subjective speech intelligibility is close to 50%. We observe that even when the noise power is higher than the attended speech power, and speech intelligibility has degraded, auditory attention decoding is still possible. Both background noise level and angular separation between competing talkers have a significant effect on speech intelligibility as well as on the neural representation of the competing speech streams.

### D. Road to neuro-steered hearing devices

For attention decoding algorithms to be incorporated in neuro-steered hearing prostheses, this study is a step towards better understanding the limitations posed by acoustic conditions on decoding performance. In this study we used the clean speech envelopes, which in a real-life scenario will not be available, for decoding. Many algorithms have been proposed to cope with realistic microphone signals (Aroudi et al., 2018; Das et al., 2017; O’Sullivan et al., 2017; Van Eyndhoven et al., 2017), where it was found that the attention decoding performance was slightly worse than with clean speech envelopes. Therefore, the current results should be considered optimistic in this respect. In addition, we used simple spatio-temporal linear decoders in order to reconstruct the attended speech envelope. Newer approaches to decoding such as neural-network (Taillez et al., 2017; Wong et al., 2018a), canonical correlation analysis (de Cheveigné et al., 2018) or state-space modelling based (Akram et al., 2016; Miran et al., 2018) decoders have the possibility of more robust attention decoding by which the boundary conditions of a neuro-steered system can be further improved.

## V. CONCLUSION

Auditory attention decoding has the potential to improve state-of-the-art noise suppression algorithms in hearing prostheses by incorporating information about the attention of the listener. The main goal of this paper was to analyze the impact of background noise and speaker separation, and their interaction, on the neural entrainment of the attended speech stream, and thereby the attention decoding accuracy. We compared the attention decoding accuracy for a range of SNRs and relative speaker positions. We also conducted behavioral tests to estimate speech intelligibility under these conditions.

We found an improvement in decoding performance with the inclusion of moderate background noise. It could possibly be due to improvement in the SNR of the EEG data because of the additional effort by the subjects. Decoding performance decreased with increasing noise power, and the degrading effect of the noise was stronger when the competing talkers were located closer to each other. We also found a significant correlation between attention decoding performance and speech intelligibility across conditions indicating that acoustic conditions with 100% speech intelligibility are not necessary for the operation of an attention decoding system.

## ACKNOWLEDGMENTS

The authors would like to thank Jelien Geivers, Tine Arras and Astrid Vandevelde for their help in conducting the experiments. We would also like to thank all the subjects for their participation in the study.

## Footnotes

1 Das et al. (2017) defined SNR values such that the attended speaker is treated as the signal, while the unattended speaker and the babble noise are both treated as noise. The input SNRs were computed at the hearing aid microphone with the highest input SNR value, and the output SNRs were computed from the output of the noise suppression filter.

